# Deep predictive coding accounts for emergence of complex neural response properties along the visual cortical hierarchy

**DOI:** 10.1101/2020.02.07.937292

**Authors:** S. Dora, S. M. Bohte, C.M.A. Pennartz

## Abstract

Predictive coding provides a computational paradigm for modelling perceptual processing as the construction of representations accounting for causes of sensory inputs. Here, we develop a scalable, deep predictive coding network that is trained using a Hebbian learning rule. Without *a priori* constraints that would force model neurons to respond like biological neurons, the model exhibits properties similar to those reported in experimental studies. We analyze low- and high-level properties such as orientation selectivity, object selectivity and sparseness of neuronal populations in the model. As reported experimentally, image selectivity increases systematically across ascending areas in the model hierarchy. A further emergent network property is that representations for different object classes become more distinguishable from lower to higher areas. Thus, deep predictive coding networks can be effectively trained using biologically plausible principles and exhibit emergent properties that have been experimentally identified along the visual cortical hierarchy.

**Significance Statement:** Understanding brain mechanisms of perception requires a computational approach based on neurobiological principles. Many deep learning architectures are trained by supervised learning from large sets of labeled data, whereas biological brains must learn from unlabeled sensory inputs. We developed a Predictive Coding methodology for building scalable networks that mimic deep sensory cortical hierarchies, perform inference on the causes of sensory inputs and are trained by unsupervised, Hebbian learning. The network models are well-behaved in that they faithfully reproduce visual images based on high-level, latent representations. When ascending the sensory hierarchy, we find increasing image selectivity, sparseness and generalizability for object classification. These models show how a complex neuronal phenomenology emerges from biologically plausible, deep networks for unsupervised perceptual representation.

## Introduction

According to classical neurophysiology, perception is thought to be based on sensory neurons which extract knowledge from the world by detecting objects and features, and report these to the motor apparatus for behavioral responding (Barlow, 1953; Lettvin et al., 1959; Riesenhuber and Poggio, 1999). This doctrine is radically modified by the proposal that percepts of objects and their features are representations constructed by the brain in attempting to account for the causes of the sensory inputs it receives (Friston, 2005; Gregory, 1980; Helmholtz, 1867; Helmholtz and Southall, 2005; Kant, 1998; Mumford, 1992; Pennartz, 2015). This constructivist view is supported, amongst others, by the perceptual psychology of illusions (Gregory, 1980; Grosof et al., 1993), but also by the uniform nature of action potentials conveying sensory information to the brain, unlabeled in terms of peripheral origin or modality (Pennartz, 2015, 2009). A promising computational paradigm for generating internal world models is predictive coding (Dayan, Hinton, Neal, & Zemel, 1995; Friston, 2005; Rao & Ballard, 1999; Srinivasan, Laughlin, & Dubs, 1982; cf. Lee & Mumford, 2003). Predictive coding posits that higher areas of a sensory cortical hierarchy generate predictions about the causes of the sensory inputs they receive, and transmit these predictions via feedback projections to lower areas, which compute the errors between predictions and actual sensory input. These errors are transmitted to higher areas via feedforward projections and are used both for updating the inferential representations of causes and for learning by modifications of synaptic weights (Rao and Ballard, 1999).

In addition to being aligned with the feedforward and feedback architecture of sensory cortical hierarchies (Felleman and Van Essen, 1991; Markov et al., 2014), the occurrence of some form of predictive coding in the brain is supported by accumulating experimental evidence. Neurons in the superficial layers of area V1 in mice navigating in virtual reality were shown to code error signals when visual inputs were not matched by concurrent motor predictions (Keller et al., 2012; Keller and Mrsic-Flogel, 2018; Leinweber et al., 2017). As expected for predictive coding, indications for a bottom-up/top-down loop structure with retinotopic matching were found by Marques et. al., 2018 for a lower (V1) and higher (LM) area in the mouse brain. In monkeys, evidence for coding of predictions and errors has been reported for the face-processing area ML (Schwiedrzik and Freiwald, 2017). In humans, predictive coding is supported by reports of spatially occluded scene information in V1 (Smith and Muckli, 2010) and suppressed sensory responses to predictable stimuli along the visual hierarchy (Richter et al., 2018).

While foundational work has been done in the computational modeling of predictive coding, there is a strong need to investigate how these early models - which were often hand-crafted and limited to only one or two processing layers (Rao and Ballard, 1999; Spratling, 2012a, 2008; Wacongne et al., 2012) - can be expanded to larger and deeper networks in a neurobiologically plausible manner. For instance, previous models studying attentional modulation or genesis of low-level response properties of V1 neurons (such as orientation selectivity) were limited to only a few units (Spratling, 2008) or to one processing layer devoid of top-down input (Spratling, 2010; Wacongne et al., 2012).

Thus we set out, first, to develop a class of predictive coding models guided by computational principles that allow architectures to be extended to many layers (i.e. hierarchically stacked brain areas) with essentially arbitrarily large numbers of neurons and synapses. Second, learning in these models was required to be based on neurobiological principles, which led us to use unsupervised, Hebbian learning instead of back-propagation (Rumelhart et al., 1986) or other AI training methods (Lillicrap et al., 2016; Salimans et al., 2017) incompatible with physiological principles.

Third, we aimed to investigate which neuronal response properties evolve emergently in both low and high-level areas, i.e. without being explicitly imposed *a priori* by network design constraints. We paid attention to both low-level visual cortical properties such as orientation selectivity (Hubel and Wiesel, 1961) as well as high-level properties such as selectivity for whole images or objects found in e.g. inferotemporal cortex (Desimone et al., 1984; Gross et al., 1972; Perrett et al., 1985).

## Materials & Methods

### Architecture of the Model with Receptive Fields

It is known that Receptive Field (RF) size increases from low to high-level areas in the ventral stream (V1, V2, V4 and inferotemporal cortex (IT)) of the visual system (Kobatake and Tanaka, 1994). To incorporate this characteristic, neurons in the lowermost area of our network (e.g. V1) respond to a small region of visual space. Similarly, neurons in the next area (e.g. secondary visual cortex (V2)) are recurrently connected to a small number of neurons in V1 so that their small RFs jointly represent the larger RF of a V2 neuron. This architectural property is used in all areas of the network, resulting in a model with increasing RF size from lower-level to higher-level areas. Furthermore, there can be multiple neurons in each area having identical RFs (i.e., neurons that respond to the same region in visual space). This property is commonly associated with neurons within cortical microcolumns (Jones, 2000).

The model variants described in this paper receive natural images in RGB color model as sensory input of which the size is described by two dimensions representing the height and width of an image. Similarly, RFs of neurons in visual cortical areas extend horizontally as well as vertically. To simplify the explanation below, we will assume that the input to the network is one-dimensional and correspondingly neurons in the model also have receptive fields that can be expressed using a single dimension. Later, we will extend the description to two-dimensional sensory input.

Figure 1 shows the architecture of the network. Consider a network with (*N* + 1) layers which are numbered from 0 to *N*. The layers 1 to *N* in the network correspond to visual cortical areas; layer 1 represents the lowest area (e.g. primary visual cortex (V1)) and layer *N* the highest cortical area (e.g. area IT). Layer 0 presents sensory inputs to the network. Below, we will use the term “area” to refer to a distinct layer in the model in line with the correspondence highlighted above. Each area is recurrently connected to the area below it. Information propagating from a lower-level to a higher-level area constitutes feedforward flow of information (also termed bottom-up input) and feedback (also known as top-down input) comprises information propagating in the other direction. Conventionally, the term “receptive field” of a neuron describes a group of neurons that send afferent projections to this neuron. In other words, a receptive field characterizes the direction of connectivity between a group of neurons and a “reference” neuron. Here, the term receptive field is used to characterize the hierarchical location of a group of neurons with respect to a reference neuron. Specifically, the receptive field of a neuron represents a group of neurons in a lower-level area that are recurrently connected to the higher-level neuron *x*. Similarly, the group of cells that receive projections from a given neuron represents the projective field of that neuron. In the current paper the term “projective field” of a neuron *x* describes a group of higher-level neurons that are recurrently connected to the lower-level neuron *x*.

**Figure 1.**
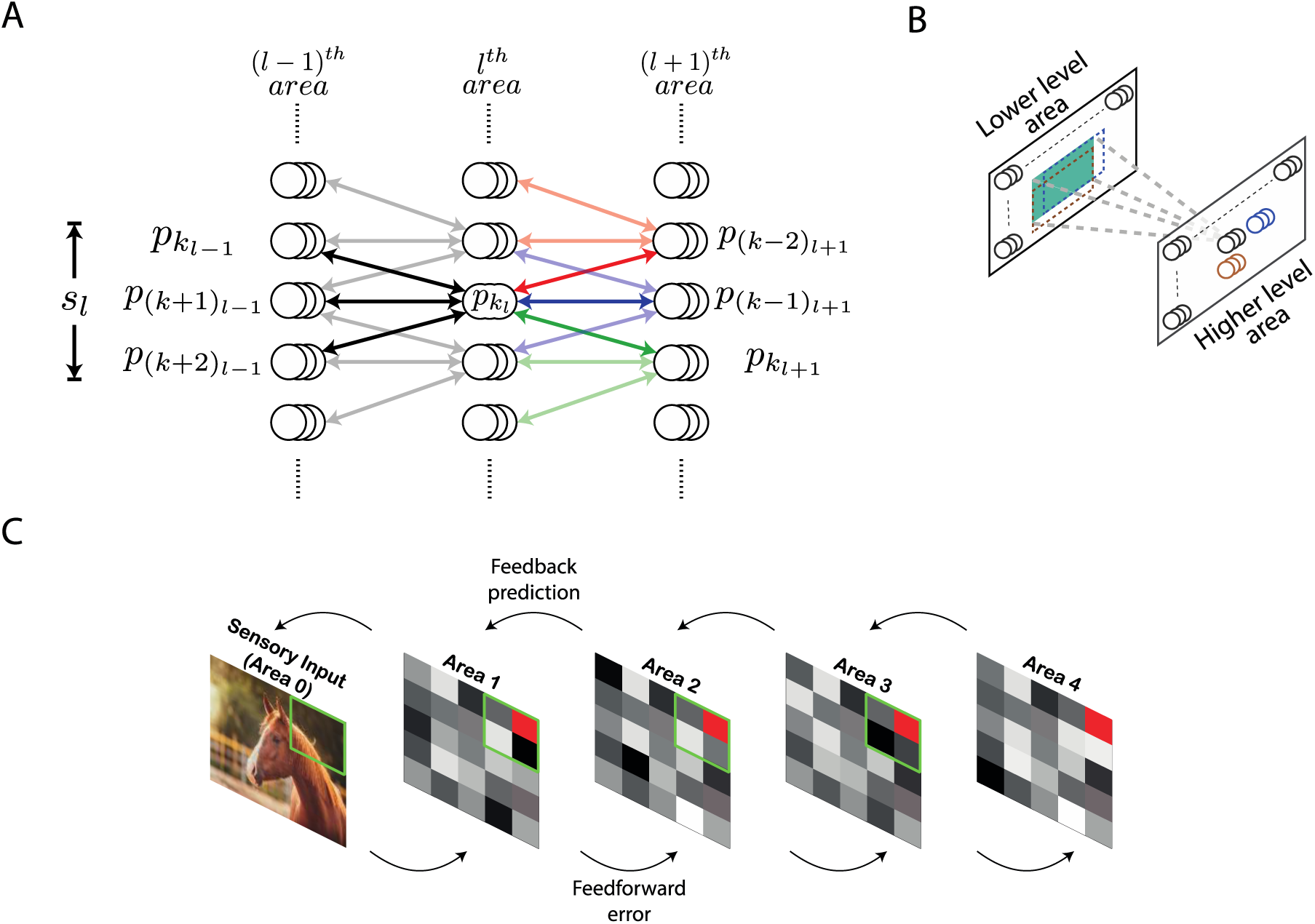
Architecture of the deep predictive coding network with receptive fields. (A) A population of neurons having identical receptive fields is represented by three overlapping circles. *p_kl_* denotes the *k^th^* population in the *l^th^* area and *s_l_* is the size of the receptive field of all populations in the *l^th^* area. Both *s_l_* and *s_l_*_+1_ have been set to 3 here. For this value of *s_l_*, the populations *p_kl_*_−1_ through *p*_(*k*+2)_*_l_*_−1_ constitute the receptive field of the population *p_kl_* (their connections are represented by black lines). Similarly, for this value of *s_l_*_+1_, *p_kl_* will be present in the projective fields of populations *p*_(*k*−2)*l*+1_ through *p_kl_*_+1_. The populations within the projective fields of *p*_(*k*−2)*l*+1_, *p*_(*k*−1)*l*+1_ and *p_kl_*_+1_ have been shown using red, blue and green arrows, respectively. Their connections with *p_kl_* are rendered in full color while other connections are shown in light colors. (B) For processing images, neuronal populations in each area can be visualized in a two-dimensional grid. Each population exhibits a two-dimensional receptive field (the receptive field of an example population in a higher-level area is shown in green). As a result, the receptive fields of two different populations can exhibit different overlaps horizontally and vertically. The receptive fields of two horizontally adjacent populations (black and blue) overlap completely in the vertical direction and partially in the horizontal direction. Similarly, the receptive fields of two vertically adjacent populations (black and brown) overlap completely in the horizontal direction and partially in the vertical direction. (C) An overview of the network with *n_l_* = 1 for all areas. Sensory input is presented to the network through Area 0. Activity of neurons in areas 1-4 is represented by tiles in grayscale colors. The green square in a lower area denotes the receptive field of the population represented as a red tile in the higher area.

Neurons in the *l^th^* area are organized in populations of *n_l_* neurons having identical receptive and projective fields. Populations having an equal number of neurons are used to reduce computational overhead. The activity of the *k^th^* population in the *l^th^* area, referred to as *p_kl_*, is a (*n_l_ by* 1) vector denoted by **y_kl_**. To reduce computational complexity, we assume that the receptive fields of all neurons in the *l^th^* area are of equal size, denoted by *s_l_*, and the receptive fields of two consecutive populations have an overlap of (*s_l_* − 1). The population *p_kl_* is reciprocally connected with populations *p_kl_*_−1_ through *p*(*k*+*s_l_*−1)*_l_*_−1_ (Figure 1). Thus, the number of populations (with distinct receptive fields) in the *l^th^* area is (*s_l_* − 1) less than the number of populations in the (*l* − 1)*^th^* area. The synaptic strengths of connections between the populations *p_kl_* and *p_kl_*_-1_ is a (*n_l_*_−1_ by *n_l_*) matrix denoted by ***W_kl-1kl_***. We assume that the neuronal populations *p_kl_* and *p_kl_*_-1_ are connected by symmetric weights, i.e. feedforward and feedback projections between these populations have equal synaptic strengths. The top-down information transmitted by population *p_kl_* to *p_kl_*_−1_ is denoted by 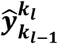 and is given by

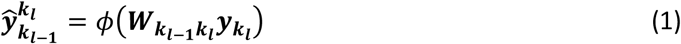

where *ɸ* is the activation function of a neuron. Predictions (see section “Learning and inference rule”) about activities of the population *p_kl_*_-1_ are denoted by 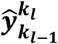. Neuronal activity is described in terms of firing rate, which by definition can never be negative. Therefore, we used a Rectified Linear Unit (ReLU) as an activation function which is defined as

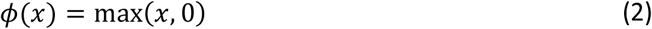

which results in values that are positive or zero. To extend the architecture described above for handling natural images, the populations in each area can be visualized as a two-dimensional grid (Figure 1B). Here, each population has receptive fields that extend both horizontally as well as vertically.

### Learning and Inference Rule

The learning rule presented in this section is inspired by the approach of predictive coding in (Rao and Ballard, 1999). Each area of the model infers causes that are used to generate predictions about causes inferred at the level below. These predictions are sent by a higher-level area to a lower-level area via feedback connections. The lower-level area computes an error in the received predictions, as compared to its bottom-up input, and transmits this error to the higher-level area via feedforward pathways. The information received by an area is used to infer better causes, which is termed the *inference* step of predictive coding, and also to build the brain’s internal model of the external environment, which is termed the *learning* step.

Figure 2 shows a possible neural implementation of predictive coding for a one-dimensional sensory input. For a given sensory input, the neuronal activities ([**y_1l_**, …, **y_kl_**, …]) of all neurons in the *l^th^* area collectively denote the causes of the sensory input inferred in this area. Based on these causes, the prediction of causes inferred in the (*l* − 1)*^th^* area is estimated according to Equation 1. Note that a given neuronal population in the *l^th^* area will generate predictions only about the neuronal populations within its receptive field (Figure 2). Therefore, neuronal populations in the *l^th^* area receive bottom-up errors via feedforward connections only from lower-level populations within their receptive field. Relative to area *l*, the bottom-up error 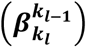 based on the prediction generated by *p_kl_* about the activity of *p_kl_*_−1_ is computed as:

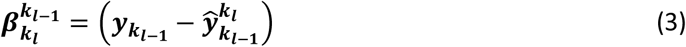

**Figure 2.**
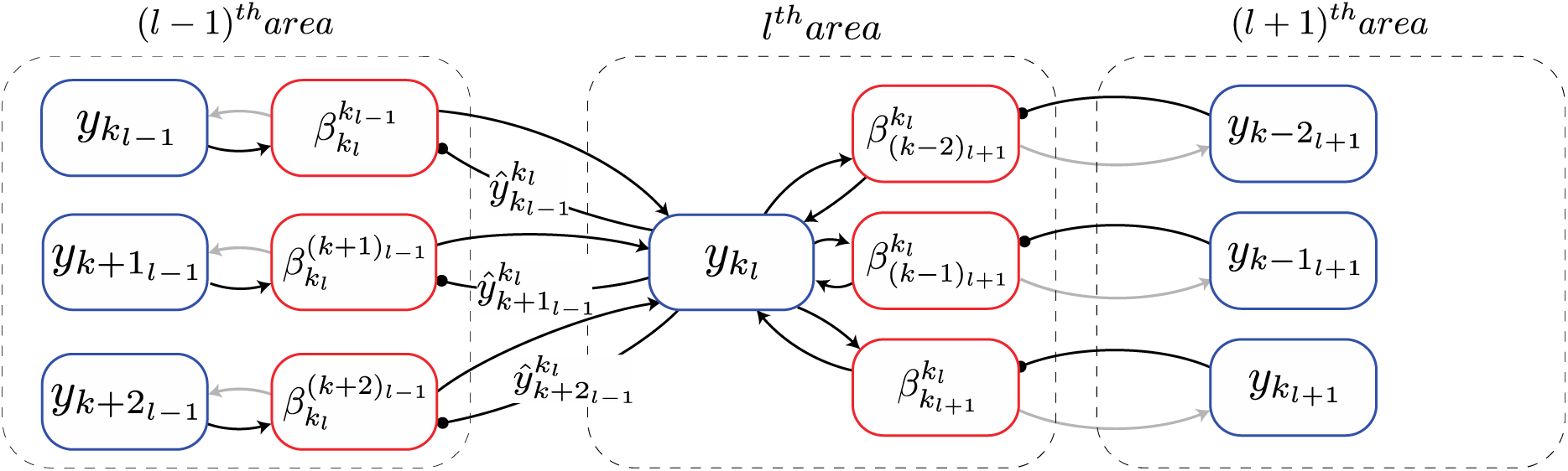
Biologically motivated realization of deep predictive coding. Each rectangle denotes a population of neurons that represents a specific signal, computed in predictive coding. The particular signal is denoted by the text inside the circle. The populations that compute errors are denoted by red blocks and the populations that represent inferred causes are denoted by blue blocks. Arrows represent excitatory connections and circles denote inhibitory connections (note that inhibitory interneurons were not explicitly modelled here). The connections that are conveying information that is required for the inference and learning steps of predictive coding are shown as black lines and other connections are shown in grey. See main text for explanation of symbols.

The computation of this bottom-up error occurs in the (*l* − 1)*^th^* area (Figure 2) and is transmitted to the *l^th^* area via feedforward projections. The simulations in this paper use a summation of squared bottom-up errors 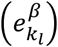 received from populations in the receptive fields of *p_kl_*, which is given as

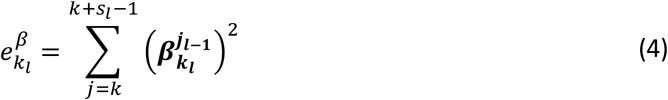

In general, other biologically plausible functions of bottom-up errors can also be used in simulations. Along with bottom-up errors, neurons in the *l^th^* area also receive a top-down prediction from neurons in the (*l* + 1)*^th^* area. Due to an overlap of (*s_l_*_+1_ − 1) between two consecutive receptive fields in area (*l* + 1), populations in the *l^th^* area will be present in the projective fields of *s_l_*_+1_ populations in the (*l* + 1)*^th^* area (Figure 1A). Populations in the *l^th^* area whose receptive fields are closer to the boundary of the visual space are an exception to this property as these neurons will be present in the projective fields of fewer than *s_l_*_+1_ populations. Here, we will focus on the general case. The population *p_kl_* will receive top-down predictions from neuronal populations *p*_(*k−sl*+1+1)*l*+1_ through *p_kl_*_+1_. The error based on the top-down prediction of the neuronal activity of the population *p_kl_* generated by the population *p_kl_*_+1_ is computed as

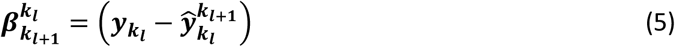

The computation of this top-down error occurs in the *l^th^* area (Figure 2). In turn, this error will also constitute the bottom-up error for the population *p_kl_*_+1_. Thus, whether an error signal is labeled bottom-up or top-down is defined relative to the area under scrutiny. The superscript and subscript in 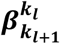 do not indicate a direction of signal propagation. The summation of squared errors due to the top-down predictions received by *p_kl_* from *p*_(*k* – *sl*+1+1)*l*+1_ through *p_kl_*_+1_ is denoted by 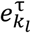 and is given as

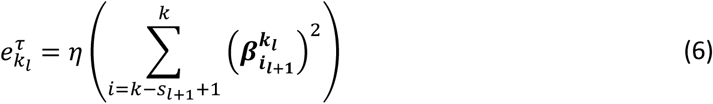

where *η* was set to one for all models unless specified otherwise (see Discussion). In addition, we employ *L*1-regularization to prevent high neuronal activities. The error due to regularization (which is symbolized by *ρ*) is given as:

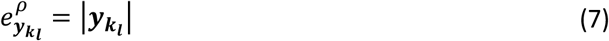

The neuronal activity of a given population is estimated by performing gradient descent on the sum of errors computed in Equations 4, 6 and 7. This results in the following update rule for inferred causes:

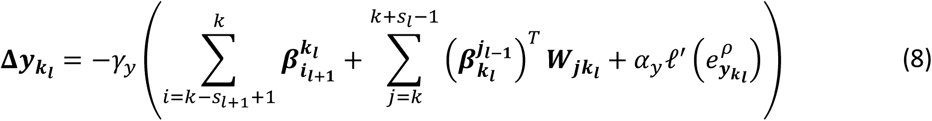

where *γ_y_* denotes the update rate for neuronal activities and *α_y_* denotes the factor which controls how strongly the regularization penalty is imposed in comparison to other errors. *ℓ*′(·) denotes the partial derivative of the regularization term. The update rule in Equation 8 constitutes the inference step of predictive coding. It results in causes that better match with top-down predictions and result in lower bottom-up errors. Higher-level areas thus influence the representations inferred in lower-level areas through top-down predictions. Similarly, lower-level areas affect the representations inferred in higher-level areas via bottom-up errors. To ensure that the neuronal activities do not become negative after updating, we rectify the neuronal activities after every inference step using the rectifier function (Equation 2). Note that **Δy_kl_** depends on the activities of neuronal populations that represent errors in the (*l* − 1)*^th^* and *l^th^* areas and the synaptic strengths of the projections between populations in these two areas (Figure 2). All of this information is available locally to the population *p_kl_*.

Moreover, the strengths of the synapses between populations in any two areas are also updated using gradient descent. As described above, an *L*1-regularization is imposed to avoid indiscriminately high values of synaptic strengths. The error due to this regularization is given as:

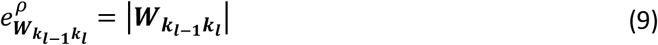

Based on the errors defined in Equations 4 and 9, the update rule for the synaptic strengths is given by

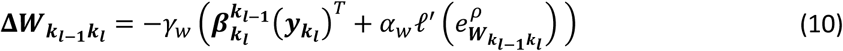

where *γ_w_* denotes the learning rate (governing synaptic weight changes) and *α_w_* is the factor which determines how strongly regularization is imposed relative to other errors. The learning rule of Equation 10 constitutes the learning step of predictive coding. Note that **Δ*W****_kl−1kl_* depends on the activity of the population that represents bottom-up errors and the activity of *p_kl_* and that these two groups are postsynaptic and presynaptic relative to each other, respectively (Figure 2). In this regard, the learning rule in Equation 10 conforms to Hebbian plasticity.

### Architecture of the Model without Receptive Fields

In the generative model described above, the inferred representations are optimized to generate an accurate prediction about causes inferred in the area below. In turn, this prediction can be used to generate a prediction about causes inferred at the next lower level. This process can be repeated until a prediction is generated about the sensory input in the lowest area. Using this method, it is possible to obtain a reconstruction of the sensory input using representations inferred in any area of the model. This functionality is shared with autoencoders (Hinton and Zemel, 1994). Here we use these reconstructions to qualitatively study the fidelity with which information about the sensory input is preserved in different areas. Our main goal is to study neural response properties in a cortex-like architecture with feedforward and feedback processing between areas, which deviates from the structure of autoencoders. Due to presence of overlapping receptive fields, neurons in each area generate multiple reconstructions of a single sensory input at the lowest level. This makes it harder to compare the reconstructions obtained using representations inferred in different areas of the model. To avert this problem, we built a network without receptive fields that is trained by the same method used for the network with receptive fields. In the network without receptive fields, each neuron in a given area is recurrently connected to each neuron in the areas below and above it. This fully connected network contained the same number of layers as the network with receptive fields and corresponding layers of the two networks contained equal number of neurons. A single reconstruction of each sensory input was obtained using the representations inferred in different areas of the network without RFs. Examples of these reconstructions are shown in the section “Reconstruction of sensory inputs”. Besides the reconstructed sensory inputs, all other results reported here are based on the results obtained with the network having RFs.

### Details of Training

The model was trained using 2000 images of airplanes and automobiles as sensory input and these were taken from the CIFAR-10 dataset. Each image has a height and width of 32 pixels. Table 1 shows the values of different hyperparameters associated with the architecture and learning rule. During training, stimuli were presented to the network in batches of 100. For each stimulus in a batch, the *inference* step (Equation 8) was executed 20 times in parallel in all areas and then the *learning* step (Equation 10) was executed once. Biologically, this corresponds to inferring representations of a sensory input on a faster time scale and updating the synapses of the underlying model on a longer time scale. At the beginning of training, the activity of all neurons in the network was initialized to 0.1 and the model was trained for 25000 iterations.

**Table 1.**
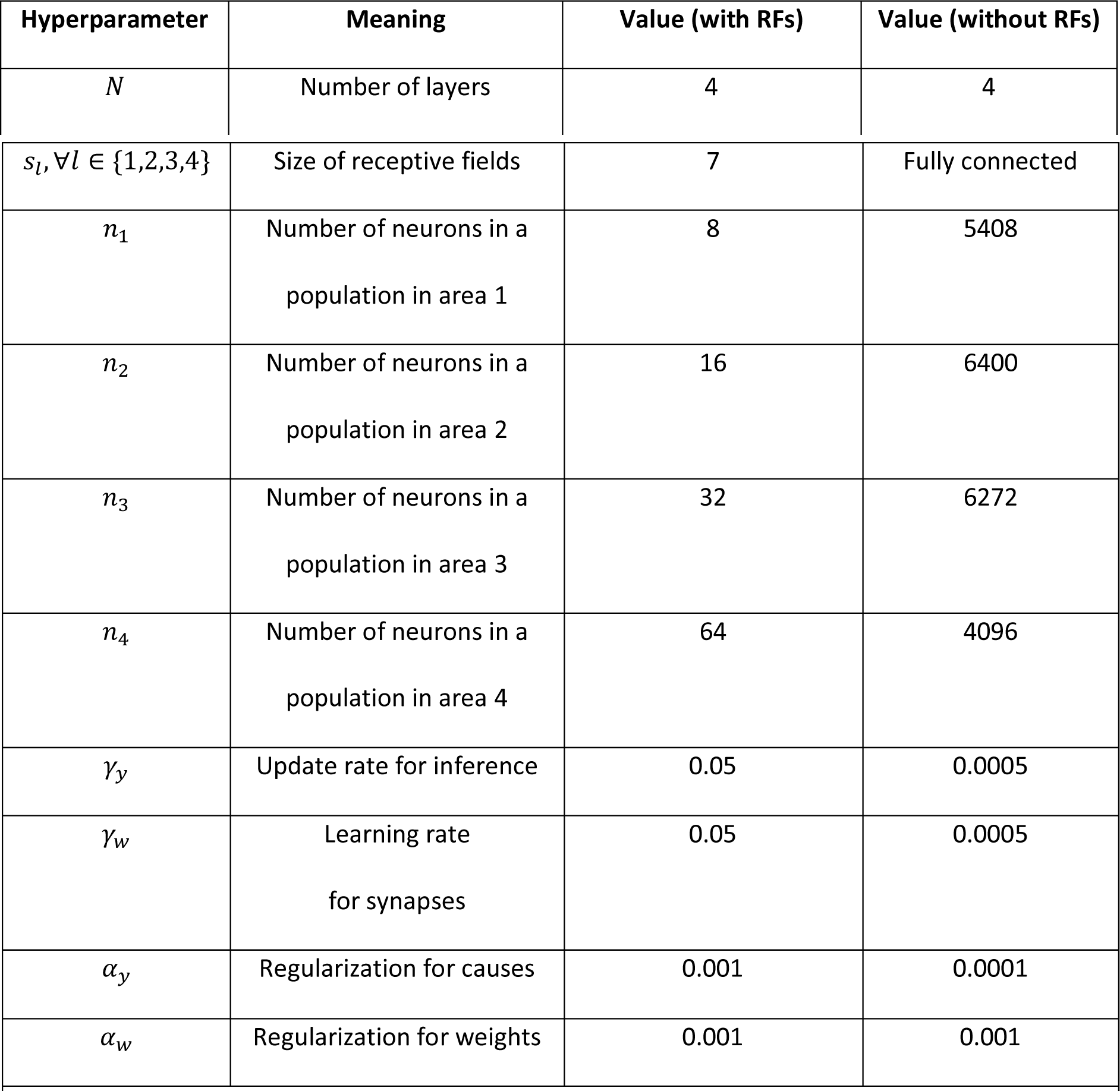
Hyperparameter settings used for training the network with and without receptive fields The size of receptive field in the network with receptive fields is equal in both image dimensions. Note that the term receptive field (RF) has been used in this table in line with its conventional definition. For the network without RFs, *n*_1_, *n*_2_, *n*_3_ and *n*_4_ are equal to the total number of neurons in each area.

Because the visual input is of equal height and width, populations in areas 1 to 4 can be visualized in a two-dimensional square grid of, for instance, sizes 26, 20, 14 and 8, respectively. Thus, areas 1 to 4 consist of 676, 400, 196 and 64 populations, respectively resulting in a total of 5408, 6400, 6272 and 4096 neurons (number of populations times population size), respectively. However, due to regularization and the rectification of causes after the inference step, some of the neurons remain inactive for all sensory inputs. These neurons have been excluded from the analysis conducted in this paper, as they would not be detected by electrophysiological methods. At the end of a typical training session for a network with the neuron counts given above, 5393, 1280, 694 and 871 neurons were active in areas 1 to 4 of the network, respectively.

To compute the number of synapses in the network, note that for every feedback synapse that transmits a prediction, there is a corresponding feedforward synapse that transmits an error (Figure 1C). Thus, the number of feedforward and feedback synapses in the network is equal. The number of feedback synapses from a population (neurons with identical receptive fields) is equal to the product of the population size in higher-level and lower-level areas and the receptive field size in the higher level area. For example, populations in areas 1 and 2 consist of 8 and 16 neurons (Table 1), respectively, and populations in area 2 have projective fields that extend by 7 units horizontally and vertically. This results in 6272 (7 ∗ 7 ∗ 8 ∗ 16) feedback synapses from a given population in area 2. Thus, the total number of synapses between two areas is equal to 794976 (area 0 and 1), 2508800 (area 1 and 2), 4917248 (area 2 and 3) and 6422528 (area 3 and 4; the number of populations times number of feedback synapses per population), respectively.

### Analysis of Neural Properties

Kurtosis is a statistical measure of the “tailedness” of a distribution. It is more sensitive to infrequent events in comparison to frequent events in the distribution. A commonly used definition of kurtosis, termed “excess kurtosis”, involves computing it for a given distribution with respect to the normal distribution. Under this definition, 3 (i.e., the kurtosis value of the normal distribution) is subtracted from the corresponding value of a given distribution. Given a set of observations (*x*_1_, …, *x_i_*, …, *x_N_*), excess kurtosis, henceforth referred to simply as kurtosis, is computed using the following equation:

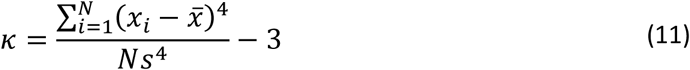

where 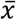 and *s* denote the mean and standard deviation of the observations (*N* in total). Based upon the use of kurtosis as a measure of neuronal selectivity (Lehky et al., 2005) and sparseness (Lehky and Sereno, 2007) in experimental neuroscience, we employ it as a measure of these properties in our model. An estimate of kurtosis obtained from responses of a single neuron to all stimuli is used as an estimate of selectivity. While computing selectivity, *N* will be equal to the number of stimuli. Similarly, its value obtained from the responses of all neurons to a single stimulus provides an estimate of sparseness. In this case, *N* will be equal to the number of neurons.

## Results

In this study we worked with two types of Deep Hebbian Predictive Coding networks (DHPC). The first type is a model without receptive fields, whereas the second model does have receptive fields. Below we will first present results from the model without receptive fields. The aim of this first modelling effort was to examine if the network is well-behaved in the sense that latent representations of causes generated in higher areas can be effectively used to regenerate the sensory input patterns in lower areas, as originally evoked by input images. Following this section we will continue with DHPC networks with receptive fields, because this type of model is better suited to examine response properties of neurons across the respective areas along the visual processing hierarchy.

### Reconstruction of sensory inputs in networks without receptive fields

For the DHPC networks without receptive fields, we used a model that was trained on an image set *X* to infer causes for an image set *Y* that was never presented to the network during training. Set *X* contains images of objects from two classes, i.e. airplanes and automobiles, and set *Y* consists of images of ten object classes namely airplanes, automobiles, birds, cats, deer, dogs, frogs, horses, ships and trucks. Note that images of airplanes and automobiles in set *Y* were different from images of these object classes in set *X*. For a given stimulus in *Y*, a separate reconstruction of this stimulus is obtained using the causes inferred from each area of the model. For a given area, the inferred causes transmit a prediction along the feedback pathways to the level below. This process is repeated throughout the hierarchy until a predicted sensory input is obtained at the lowest level. Figure 3 shows examples of reconstructions of novel stimuli obtained using the causes inferred in each area of the model, along with the original sensory input. The first three exemplars are of airplanes and an automobile which belong to object classes that were used to train the model. The other exemplars are reconstructions of a frog, a bird, a horse and a ship, which were never presented to the network during training, neither as exemplar nor as object class. We conclude that the reconstructions become somewhat blurrier if the generative process is initiated from higher, as opposed to lower, areas of the model, but also that the natural image statistics are captured reasonably well. This is remarkable because these inputs had never been presented to the network before.

**Figure 3.**
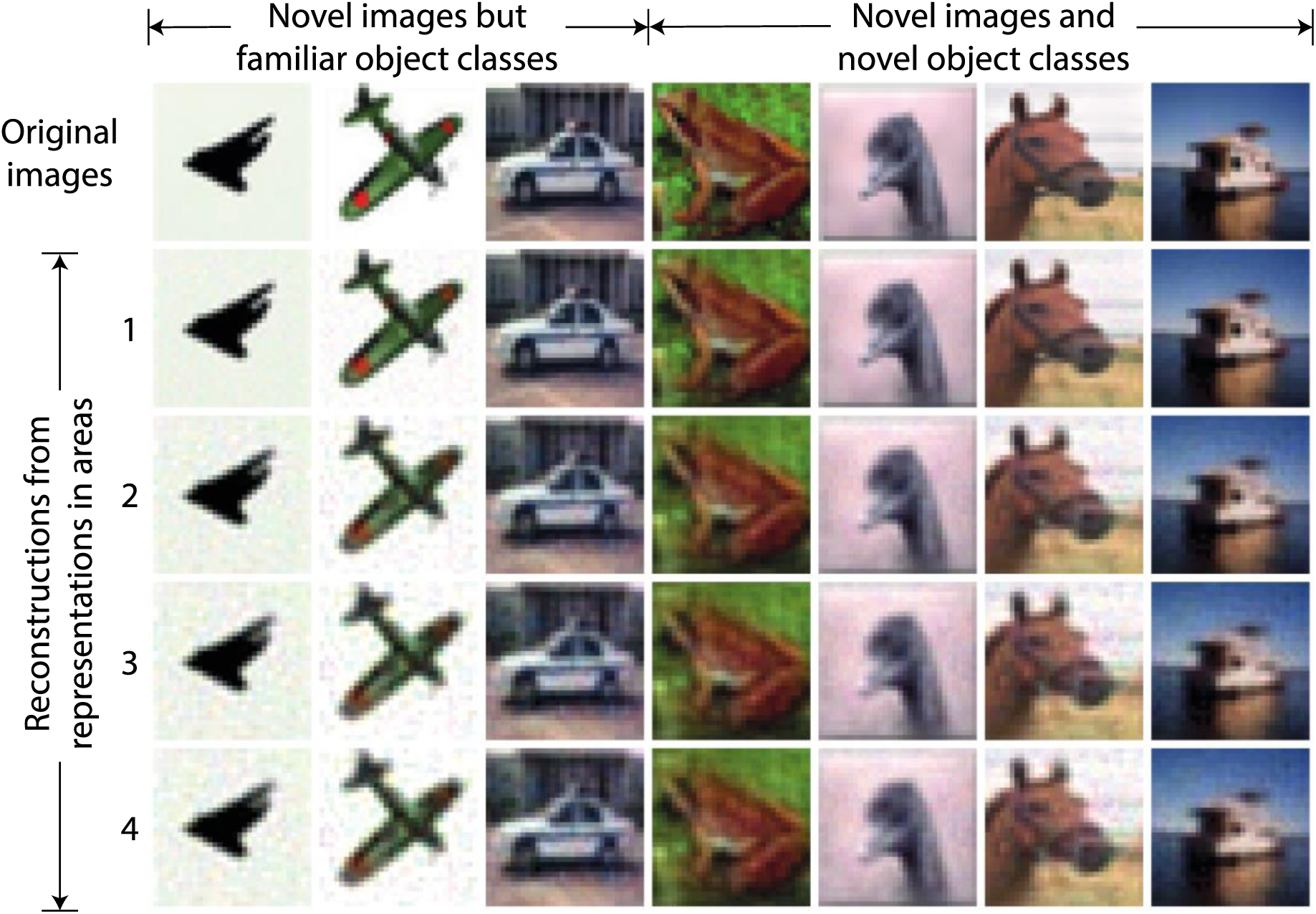
Examples of reconstructions obtained using causes inferred by the trained model without receptive fields. Each column represents an example of a sensory input. The three leftmost images represent novel stimuli from object classes used in training whereas other images are from object classes not used in training. The top row shows the novel sensory input that was presented to the network to allow it to construct latent representations across the areas. Rows 2 to 5 show the reconstructions of the sensory input obtained using the latent representations in the corresponding areas of the model. It can be observed that the reconstructed sensory input faithfully reproduces the novel originals, although the lower areas regenerate the inputs more sharply.

### Orientation selectivity in a lower area of the network with receptive fields

Neurons in V1 respond selectively to sensory input consisting of edges oriented at specific angles in their receptive fields (Hubel and Wiesel, 1959). The neurons in layer 1 of the model with receptive fields also exhibited this property. Importantly, this orientation selectivity was not hand-crafted or built into the network a priori, but emerged as a consequence of training the network on inputs conveying naturalistic image statistics. After training, the strengths of feedback synaptic connections between area 1 and 0 of the model resembled Gabor-like filters. Figure 4 shows plots of strengths of synapses onto a given neuron as representative examples for area 1 of the model (Figure 1C). These plots were obtained by normalizing the feedback weights of a representation neuron in area 1 to the interval [0, 1]. Each image is obtained by rendering the normalized weights of a single representation neuron in area 1 as pixel intensities where each pixel corresponds to a specific neuron in area 0 in the receptive field of this representation neuron. Conventionally, orientation selectivity is viewed as a property of feedforward projections to V1. The model described here uses symmetric feedforward and feedback weights (apart for their difference in sign, fig. 2), therefore the orientation selectivity illustrated here is applicable to both feedforward and feedback connections between areas 0 and 1.

**Figure 4.**
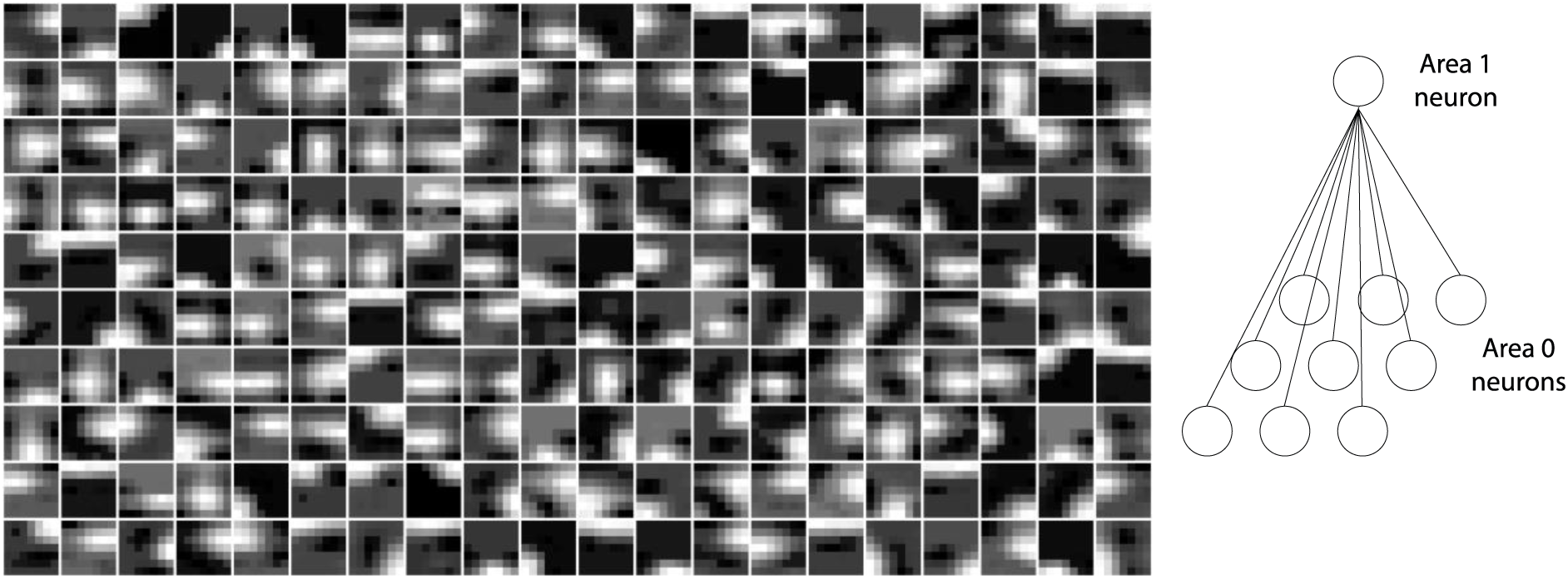
Emergence of orientation selectivity in the lowermost area (area 1) of a trained model with receptive fields. Plots show normalized synaptic strengths for connections between area 1 and 0 (i.e. the input layer) of the model. Each box shows a symbolic representation of synaptic strengths from a randomly selected area 1 neuron to all area 0 neurons within its receptive field (right panel). Darker regions in the images correspond to synaptic strengths closer to zero and brighter regions in the images correspond to strengths closer to 1. It can be observed that receptive fields of many cells contain non-isotropic patches imposing orientation selectivity on neural responses in area 1.

### Image Selectivity

Neurons in different brain areas situated along the sensory processing pathways exhibit tuning to features of increasing complexity. Whereas neurons in the primary visual cortex (V1) respond to edges of different orientations (see above) neurons in V4 respond selectively to e.g. textures and colors (Okazawa et al., 2015) and neurons in IT show selectivity to particular faces or other objects (Gross et al., 1972; Logothetis and Pauls, 1995; Perrett et al., 1992; Tanaka et al., 1991). This property is manifested by differences in neuronal selectivity across areas of the visual cortical hierarchy with later stages exhibiting higher selectivity in comparison to earlier stages. For our model, we asked whether analysis of area-wise neuronal activity would also reveal increasing selectivity from the lowest to highest areas.

Figure 5 shows the distribution of image selectivity for neurons in each area of the model. The kurtosis was computed for each neuron based on its responses to all stimuli presented to the model (Equation 11) and used as a measure of image selectivity for a single neuron (Lehky et al., 2005). The figure shows that the mean image selectivity increases from the lowest to the highest area in the model. We compared the average selectivity in a given area with every other area in the model using Mann-Whitney’s U test with Bonferroni correction for multiple comparisons. For all comparisons, the null hypothesis was rejected with *p* < 5.10^−15^. Thus, image selectivity strongly increased when ascending in the visual cortical hierarchy. Importantly, this property was emergent in the sense that it was not preprogrammed in our algorithm.

**Figure 5.**
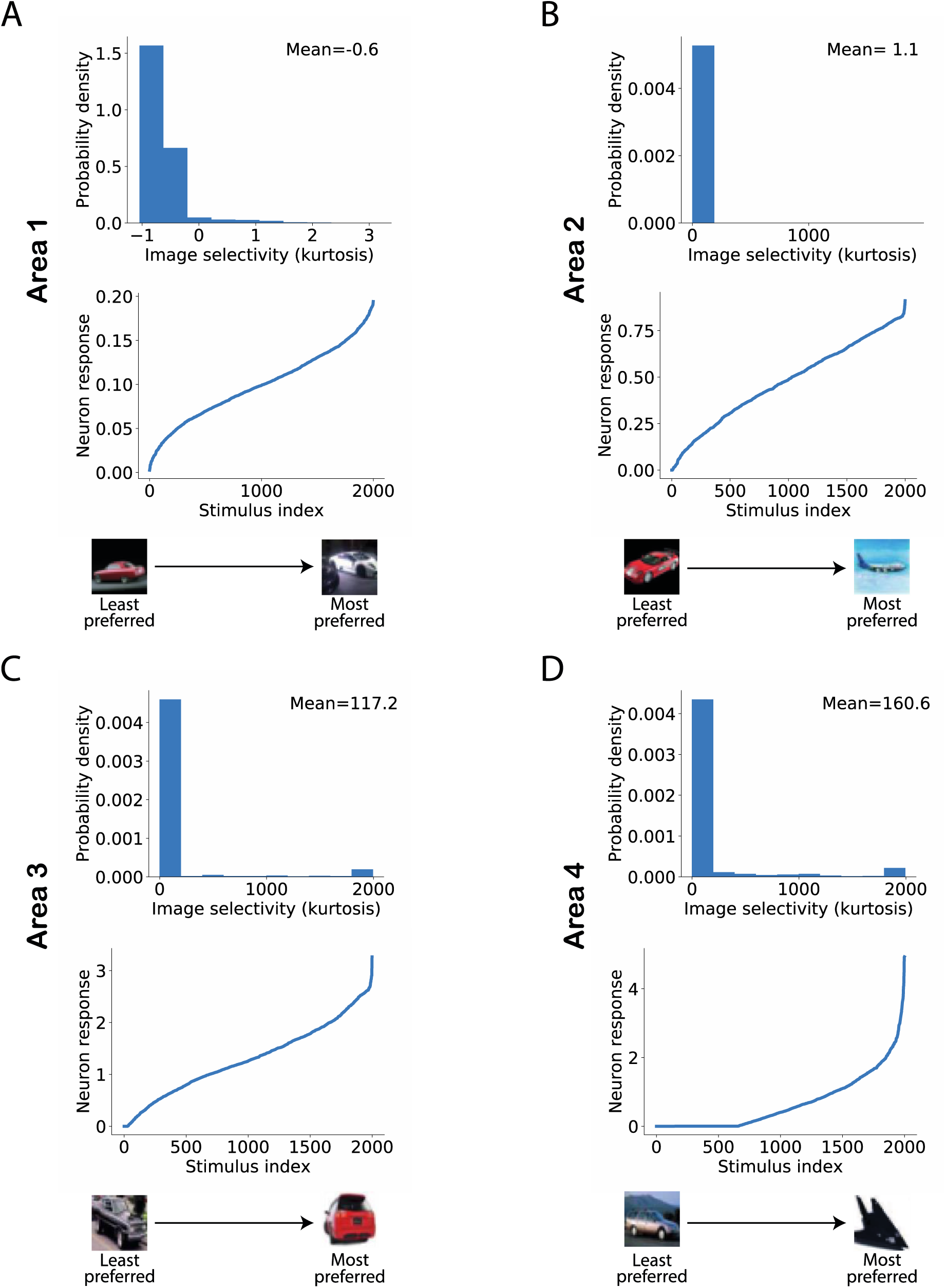
Image selectivity of model neurons. (A-D) Distribution of image selectivity of neurons in each area of the model (top panels; A: lowest area/Area 1; D: highest area/Area 4). The mean value of neuronal image selectivity for each area is shown in the top right corner of the corresponding plots. (Bottom panel) The activity of a randomly chosen neuron in each corresponding area has been sorted according to its response strength for all stimuli presented to the network. It can be observed that the average selectivity of neurons increases from lower to higher areas in line with experimental data.

### Sparseness

A feature related to neuronal selectivity is sparseness, reflecting how scarcely or redundantly a feature or object is coded across the population in a given area (Montijn et al., 2015; Perez-Orive et al., 2002; Vinje and Gallant, 2000; Willmore and Tolhurst, 2001). A high or low sparseness can easily arise in a population with large variations in average neuronal activity. For instance, consider a population in which a single neuron has an average firing rate of 100 spikes/sec and all other neurons have an average firing rate of 10 spikes/sec. In this population, the peak in the distribution of population activity due to the neuron with high average activity will result in high sparseness. To overcome this problem in the analysis, we normalized the activity of all model neurons using their average activity and an individual estimate of kurtosis was obtained for each stimulus across all neurons in each area based on this normalized activity. Figure 6 shows a distribution of sparseness in each area. We found that the average value of sparseness across all stimuli in each area increased systematically from the lowest to highest area. For validation, we conducted a pairwise comparison of sparseness values in different areas using Mann-Whitney’s U test with Bonferroni correction for multiple comparisons. For all comparisons between areas, the null hypothesis was rejected with *p* < 5.10^−34^ in all cases.

**Figure 6.**
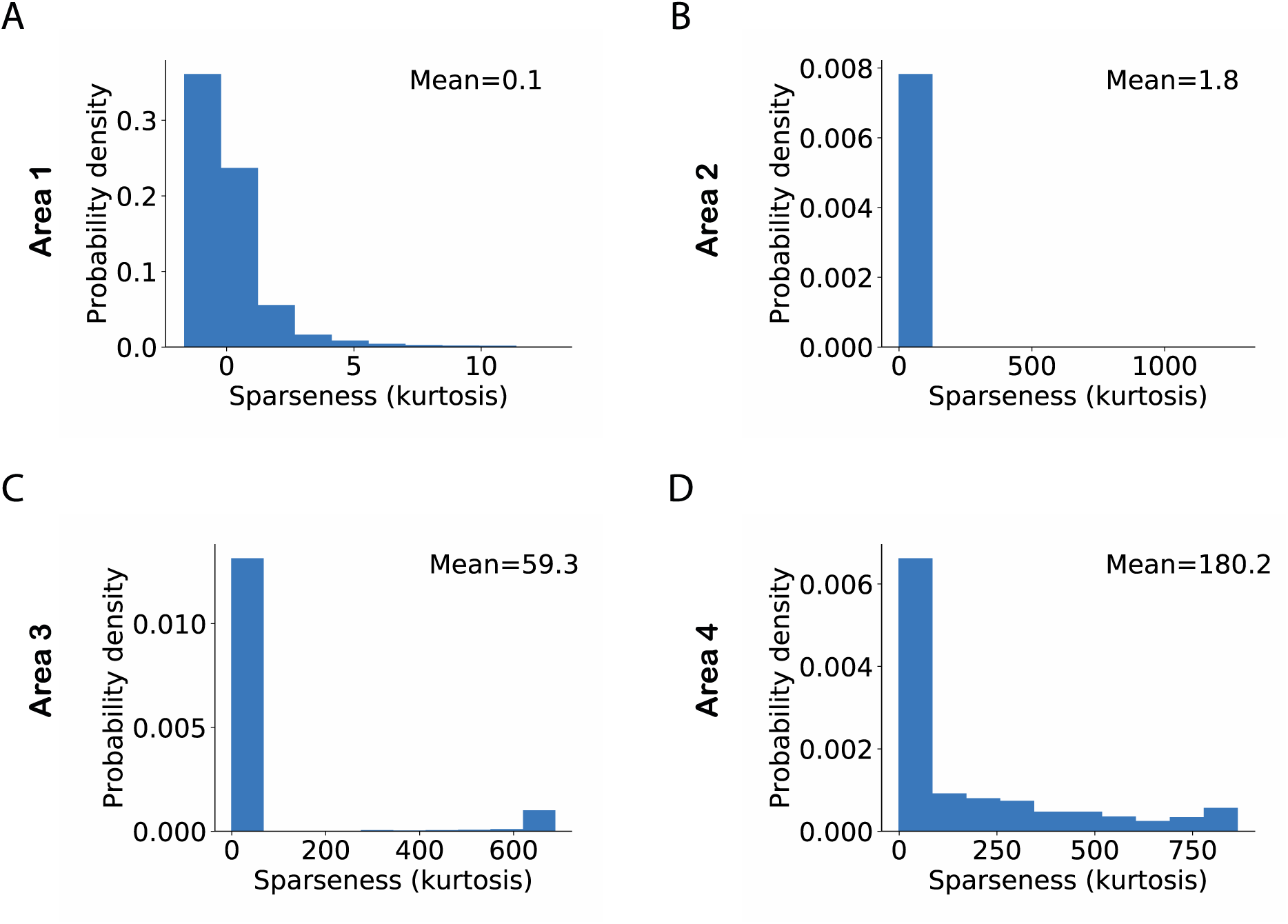
Sparseness in neuronal activity across ascending areas of the model. Sparseness was measured as the kurtosis across all neuronal responses in a given area and given a single stimulus. The mean value of sparseness is computed by averaging these estimates of kurtosis across all stimuli. (A-D) Distribution of sparseness in each area. The mean value of sparseness for each area is shown in the top right corner of each plot. It can be noted that the average sparseness of all neurons in model areas increases from lower to higher areas in agreement with some of the experimental studies.

### The relationship between the response magnitude of neurons, selectivity and sparseness

We next studied the relationship between a neuron’s average response to all stimuli and its selectivity. Similarly, for each area of the model we also investigated the relationship between a population’s average response to a stimulus and its sparseness. The selectivity in different areas of the model exhibited wide variations. For the purpose of visualizing how the relationship between selectivity and mean neuronal activity evolves from lower to higher areas, we looked at the relationship between the log of selectivity and mean neuronal activity. We observed that, in all areas, there was a negative correlation between the selectivity and average neuronal activity, i.e. neurons with high selectivity had low average activity. Pearson correlation coefficients of −0.23, − 0.05, −0.55 and −0.42 were obtained between selectivity and mean responses in areas 1 to 4, respectively. This has also been reported in experimental data (Lehky et al., 2011). Further, this negative correlation became stronger from lower to higher areas in the model.

We conducted a similar study on the relationship between sparseness and average population activity. It has been reported in experimental data that the average population response shows little variation for different values of sparseness (Lehky et al., 2011). This was also the case for all model areas as we observed only weak correlations between sparseness and average population responses. Pearson correlation coefficients of −0.18, 0.02, 0.23 and 0.18 were obtained between sparseness and mean responses in areas 1 to 4, respectively. These similarities between the statistical properties of model neurons and data from animal experiments arise without being imposed by network design or training procedure.

### Impact of neuronal selectivity and neuronal response range on sparseness

Although selectivity and sparseness represent different aspects of neuronal activity, they are interconnected quantities, i.e. a population consisting of highly selective neurons will also exhibit sparseness in the population response to a single stimulus. However, it has also been observed in data recorded from macaque IT that the dynamic range of neuronal responses correlates more strongly with sparseness than selectivity (Lehky et al., 2011). Here, dynamic range was quantified using the interquartile range of neuronal responses, which is the difference between the 75^th^ and 25^th^ percentiles of a neuron’s responses to the individual stimuli presented. We asked which of the two factors, selectivity or dynamic range, contributed to sparseness in the responses of model neurons in different areas.

To examine the interactions between these network parameters, we estimated sparseness in three different sets of neuronal populations that differed in terms of selectivity and dynamic range. Figure 7 shows the histogram of interquartile ranges for neurons in each area. It can be observed that the dynamic range gradually increased from lower to higher areas as more neurons shifted away from low range values. For each area, we considered a first subset, denoted by ‘SNR’ (i.e., Selective Neurons Removed), obtained by removing activities of the top 10% of neurons having the highest selectivity in that area. To obtain the second subset of each area, denoted by ‘DNR’ (i.e. Dynamic range Neurons Removed), we eliminated the activities of the top 10% of neurons with the broadest interquartile ranges. Figure 8 also shows the distribution of sparseness of the third set, viz. including all neurons of an area (denoted by ‘All’) as well as for the two subsets described above. It can be clearly seen that sparseness is more dependent on neurons with high selectivity in comparison to neurons that exhibit a broad dynamic range. Thus, our model shows a strong influence of neuronal selectivity on sparseness. However, this behavior of the model was dependent on regularization (see Discussion).

**Figure 7.**
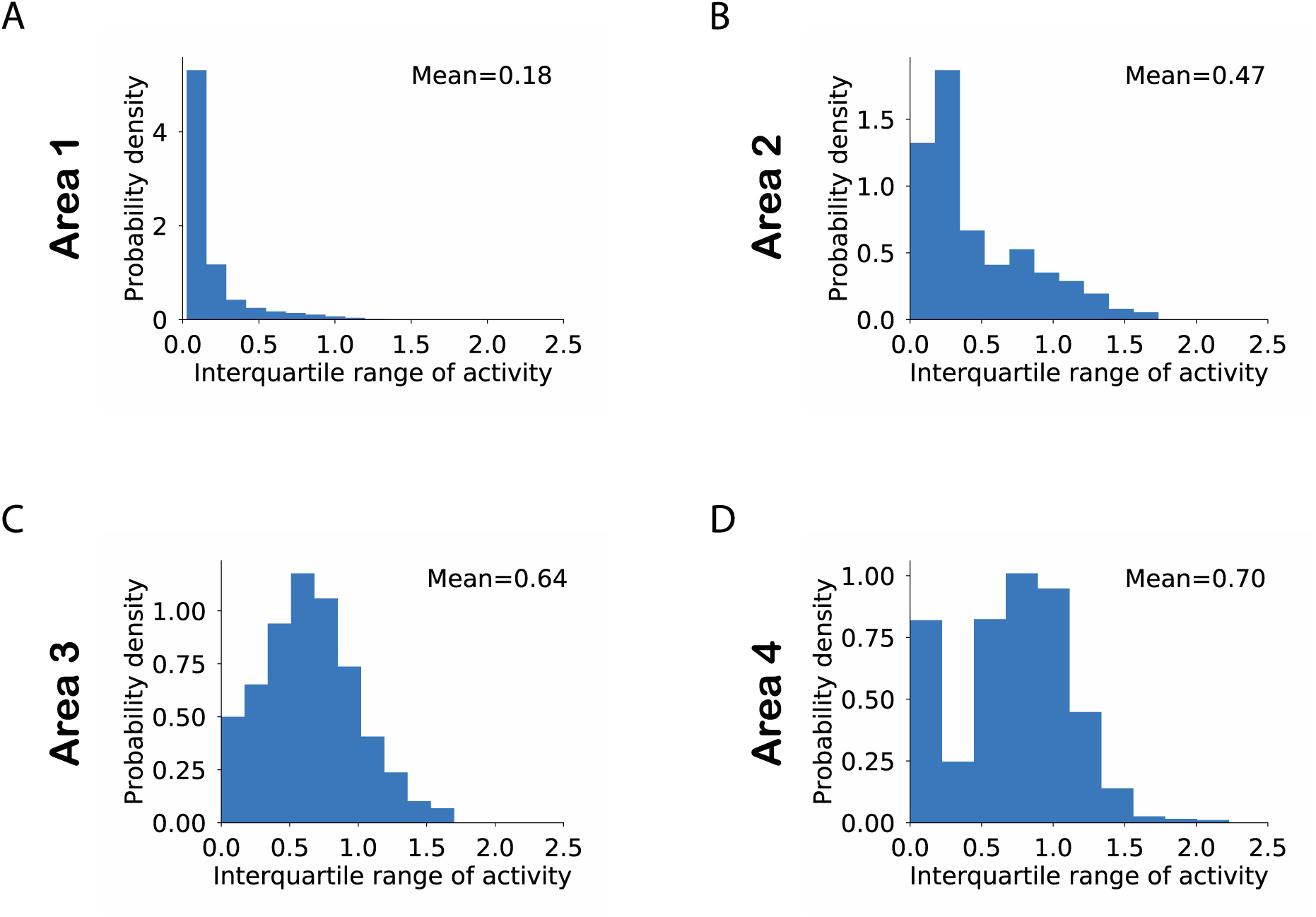
(A-D) Distribution of the dynamic range of neurons computed as the interquartile range of the neuronal responses in a given area across all stimuli. The mean value for each area is computed by averaging across interquartile ranges for all neurons in that area.

**Figure 8.**
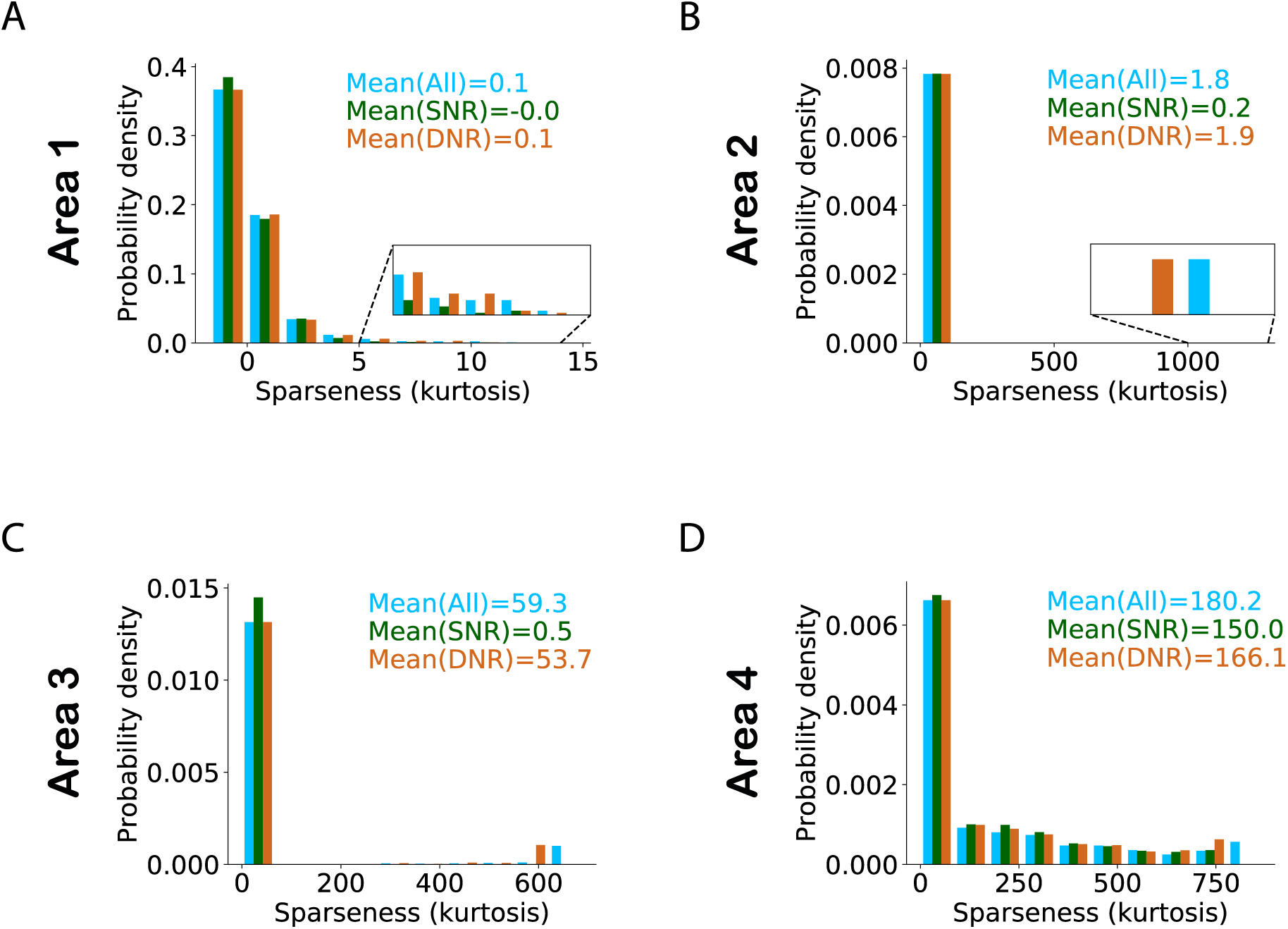
Effect of high selectivity and high dynamic response range neurons on sparseness. Histogram of sparseness for three different populations of neurons. The distribution of sparseness for all neurons is shown in blue. The population in which the top 10% most selective neurons were removed (SNR) is shown in dark green and light brown color denotes the populations in which neurons with high dynamic response range were removed (DNR). Values represent the mean sparseness estimates for the different populations in corresponding colors. In all areas of the model (except area 1) it can be observed that the mean sparseness drops much more strongly on removal of highly selective neurons in comparison to removal of neurons with high dynamic range.

### Object classification performance

We next studied the ability of the model with RFs to infer causes that generalize across different exemplars of a given object class. The exemplars varied in terms of object identity, viewing angle, size, etc. For this purpose, we trained separate Support Vector Machine (SVM) classifiers using latent representations of causes in each area of the model. Using a subset of the stimuli with which the model was trained, a linear SVM classifier was optimized to distinguish between representations of exemplars of two object classes, i.e. airplanes and automobiles. The remaining stimuli were used to estimate the performance of the SVM classifier which thus yields an estimate of the model’s capacity to generalize across different exemplars of the same class.

To examine whether the representations in different areas exhibited better generalization progressively across ascending areas, we optimized a linear SVM classifier using representations for 1500 stimuli randomly chosen from both classes and then computed its classification performance on the remaining 500 stimuli. This analysis was repeated 100 times by bootstrapping without replacement the samples selected for optimizing the linear SVM classifier. Figure 9B shows the classification performance of the SVM classifier for representations in different areas of the model. First, we observe a classification accuracy well above chance level in all areas (one sample t-test; p-values are lower than 8.10^−130^ for all areas). Second, we observed a modest but systematic increase in the classification performance from the lowest to highest area of the model. This shows that representations in higher areas can generalize better across unfamiliar exemplars than lower areas. To validate our results, we compared the accuracy in the topmost area with accuracy in other areas using Mann-Whitney’s U test with Bonferroni correction for multiple comparisons. The maximum p-value of 0.0004 was obtained for the comparison between the accuracies of the topmost area and area 2. Based on these comparisons, the null hypothesis for all comparisons between areas was rejected at a significance level of at least 0.01. The maximum p-value of 0.0004 was obtained for the comparison between the accuracies of the topmost area and area 2.

**Figure 9.**
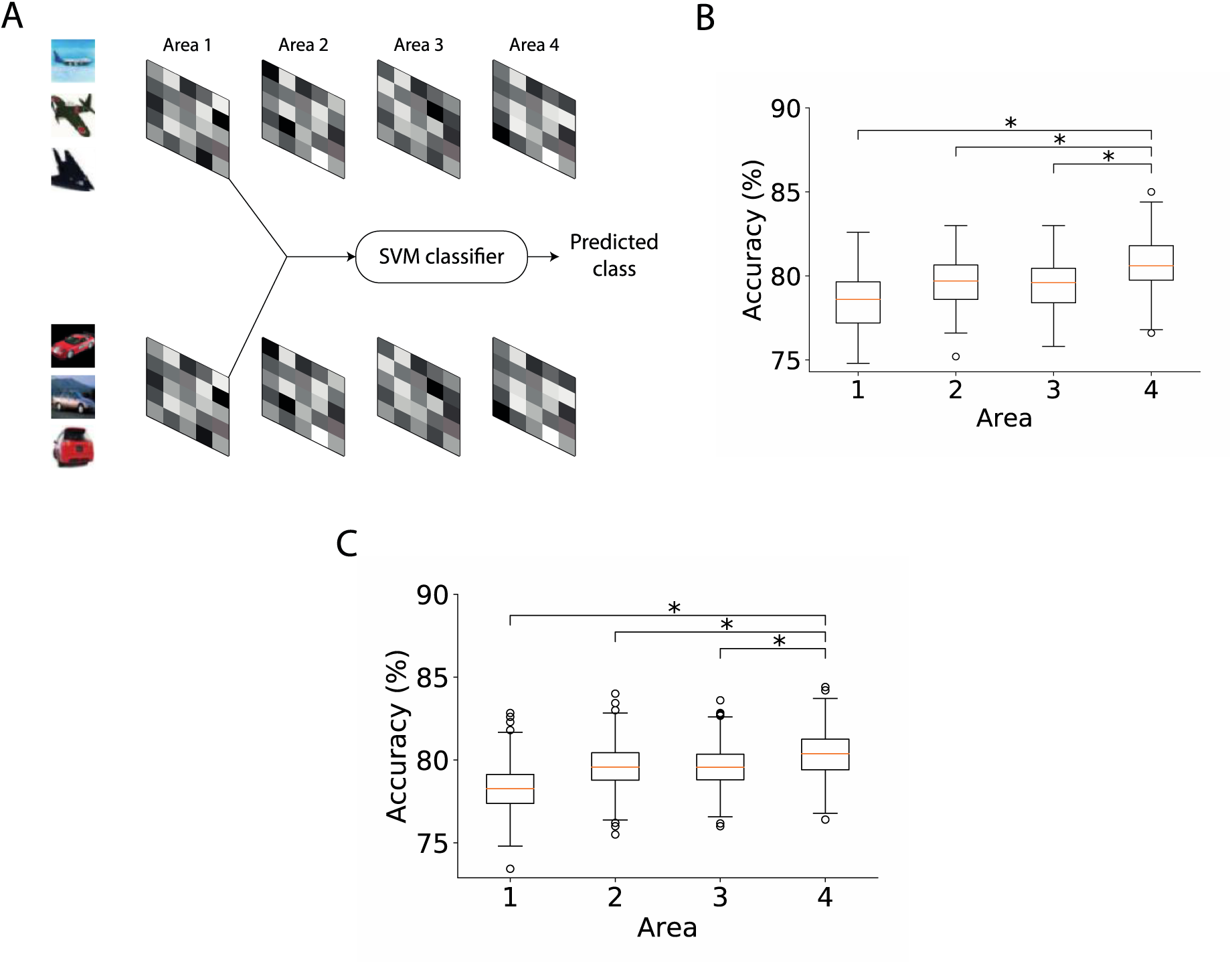
Object classification performance based on the representations of inferred causes across ascending areas. (A) Method used for computing the accuracy of a classifier based on causes, in this case, inferred in area 1. The inferred causes for a given stimulus are presented to a Support Vector Machine (SVM) classifier whose output is used to determine the predicted class (airplanes vs cars) of a given stimulus. This procedure is repeated for all areas. (B) Boxplot of classification performance in different areas using 1500 randomly selected samples for optimization. Horizontal lines of the boxes denote the first, second and third quartiles. Whiskers represent the entire range of data and circles denote outliers. The second quartile in all areas was significantly above chance level accuracy (one sample t-test, \**p* < 0.05). The performance of the classifier optimized using area 4 representations was significantly higher than the performance of classifiers of other areas (Mann-Whitney’s U test with Bonferroni correction, \**p* < 0.05). (C) Boxplot of classification performance in different areas using different numbers of samples for optimization. The number of samples did not affect the conclusions observed in (B) (Mann-Whitney’s U test with Bonferroni correction, \**p* < 0.05).

To ensure that this result was not dependent on the number of stimuli used, we repeated this analysis with different stimulus sets. For this purpose, we optimized the SVM classifier on stimulus sets containing 1000 to 1500 stimuli in steps of 100 and evaluated its performance on the remaining stimuli. Figure 9C shows the performance of the classifiers optimized using different numbers of stimuli for different areas of the model. The generalizing capacity of the inferential representations in higher areas of the model was better than in the lower areas irrespective of the number of stimuli used to optimize the SVM classifier. For all comparisons, the null hypothesis could be rejected at a significance level of at least 0.05. The lowest level of significance was obtained for the comparison between the accuracies of the top area and area 2 (*p* < 1.10^−21^). Again, this model behavior arose emergently as it was not pre-programmed or built a priori into the network design.

## Discussion

First, we described a general method to build neurobiologically plausible deep predictive coding models for estimating representations of causes of sensory information. Different hyperparameters of the network can be modified to model various aspects of cortical sensory hierarchies; for instance, *N* can be varied from 1 to 5 to study cortical hierarchies of increasing depth. This provides a mechanism to develop deep neural network models of information processing in the brain that can be used to simultaneously study properties of lower-level as well as higher-level brain areas. The models were trained using unsupervised, Hebbian learning and both the inference and learning steps utilized only locally available information. Second, we found that several properties of neuronal and population responses emerge in the model without being imposed by network design or by the inference and learning steps. Image selectivity increased systematically from lower levels to higher levels and the average sparseness of inferred representations increased from lower levels to higher levels, which is in line with at least some experimental study (Okazawa et al., 2017). Hereby DHPC networks provide a biologically plausible solution to the problem of ‘combinatorial explosion’ which would arise if the occurrence of strongly object-selective (“grandmother cell”) responses has to be explained from the combination of individual, low-level features (Barlow, 1972; Riesenhuber and Poggio, 1999).

Furthermore, we studied object classification properties of the causes inferred by the model. The classifiers optimized using representations in higher areas exhibited better performance in comparison to those using lower-area representations. Thus, predictive coding may provide a useful basis for the formation of semantic concepts in the brain, at least when combined with networks performing categorization (e.g. in the medial temporal lobe (Quiroga et al., 2005) or prefrontal cortex (Freedman et al., 2003)).

### Reproduction of experimental findings by the model

The increase in image selectivity in ascending areas of DHPC networks has also been reported in experimental studies (Gross et al., 1972; Logothetis and Pauls, 1995; Tanaka et al., 1991). This can be attributed to the property that neurons in each model area are strongly active when the neurons within their receptive field exhibit a particular pattern of activity. For example, neurons in the lowest area of the model develop Gabor-like filters that resemble oriented edges and have been shown to form a representation code for natural scenes that consists of statistically independent components (Bell and Sejnowski, 1997). These low-level neurons will be strongly active when a particularly oriented edge is present within its receptive field. Similarly, a neuron at the next level will be strongly active when neurons within its receptive field at the lower level exhibit a specific pattern of activity. This implies that a neuron at this higher level will only become active when a particular configuration of edges (rather than a single edge) occurs at a specific location in visual space, resulting in increased in complexity of features detected by neurons at this level. This increase in feature complexity of features detected by neurons in successive model areas leads to a corresponding increase in the average neuronal selectivity when ascending the hierarchy.

It could be argued that regularization will automatically lead to an increase in average selectivity in neuronal responses across model areas. To examine this possibility, we also trained models with no regularization (neither for synaptic weights nor inferred causes) while all other hyperparameters remained unchanged. These models also exhibited an increase in average selectivity across model areas (data not shown). However, adding regularization did result in an overall increase in average selectivity in each of the model areas. By definition, the responses of a selective neuron will have a high interquartile range. Thus, the increasing selectivity across model areas also leads to an increase in the average interquartile range across ascending model areas (Figure 7).

Unlike selectivity, there is no consensus in the literature on how sparseness varies along the cortical hierarchy due to a lack of consistency in experimental data. Responses of macaque V4 neurons were reported to exhibit higher sparseness in comparison to V2 neurons (Okazawa et al., 2017). In line with our results, these findings indicate that sparseness increases from lower-level to higher-level areas. In another study, however, it was shown that sparseness estimates based on responses of macaque V4 neurons did not differ significantly from estimates for IT neurons (Rust and DiCarlo, 2012). Both of the above experimental studies quantified sparseness using the same two measures, namely the sparseness index described by (Vinje and Gallant, 2000) and entropy (Lehky et al., 2005). Although sparseness was quantified here using kurtosis, its estimates across different areas of the model exhibited the same relationship with one another when Vinje and Gallant’s (Vinje and Gallant, 2000) index of sparseness was used (figure not shown).

### Regulation of sparseness

Regularization had a strong influence on both average sparseness in each model area and on the relationship between average sparseness in different model areas. In the absence of any regularization, average sparseness first increased and then decreased when ascending across areas (Figure S1). This can be attributed to the network property that all areas in the model infer causes that reconcile bottom-up and top-down information (Equation 4 and 6) received by an area, except for the top area where causes are determined only by bottom-up information. This lower constraint on the top area leads to a decrease in sparseness in areas farther away from the sensory input layer. Imposing regularization only on representations inferred in areas farther from the top to compensate for this lack of constraint did not alter this pattern of average sparseness across model areas (Figure S2). This is because sparse neuronal activity in higher areas induced by regularization results in sparse top-down predictions for lower areas which indirectly induce sparseness in representations inferred in lower areas. In this manner, sparseness induced in higher areas *spreads* throughout the network. Thus, regularization in higher areas leads to an increase in average sparseness in all model areas but does not alter the overall pattern of sparseness across different model areas. However, sparseness imposed by higher areas onto lower areas can be weakened by scaling down the errors due to top-down feedback, for example, using a value of *η* < 1 in Equation 6. Thus, sparseness depends strongly on multiple factors which include regularization, hierarchical position of an area, and the weights given to bottom-up and top-down errors. These results may provide an explanation for inconsistent results regarding sparseness observed in experimental data. In experiments, sparseness has been compared across two brain regions at most, and our model suggests that results obtained from such studies may not generalize to other brain regions.

Regularization was also a factor that affected whether high selectivity neurons or high dynamic range neurons contributed strongly towards sparseness in a given area (Figure 8). In the absence of regularization, sparseness in lower areas was determined by high selectivity neurons, but in higher areas sparseness was determined by high dynamic range neurons (Figure S3). This can be attributed to the network property that the bottom-up input to lower areas is more strongly driven by a fixed sensory input whereas in higher areas the bottom-up drive is based on constantly evolving representations. Stochastic fluctuations resulting from these evolving representations at the inference step in higher areas lead to higher dynamic response ranges in these very areas. As a result, sparseness is more strongly determined by high dynamic response range neurons in higher areas, which is in line with the experimental results of (Lehky et al., 2011). However, adding regularization to the top area in the model constrains neural activity throughout the model, thereby reducing the dynamic response range of neurons (Figure S4). Furthermore, high regularization leads to neurons that are active for a small number of images. When the activity of such neurons is normalized by their mean activity, this can result in very high (relative) activity for some of these images. An estimate of kurtosis obtained from normalized neuronal activity can thus lead to arbitrarily high estimates of sparseness (Figure 8).

The relationship between statistical properties (selectivity and sparseness) of inferred representations is loosely consistent with the idea of ergodicity in experimental data. As defined in (Lehky et al., 2005), a neural system is termed ‘*weakly* ergodic’ if the average selectivity of individual neurons across multiple stimuli is equal to the average sparseness. Experimental evidence for ergodicity has been reported in multiple cortical areas (Kadohisa et al., 2005; Verhagen et al., 2004). The average selectivity and sparseness of representations inferred by the model do not satisfy this equality but there is a close relationship between these two properties, as removal of highly selective neurons strongly degrades sparseness (Figure 8). Possibly, equality of average selectivity and sparseness is only satisfied under certain hyperparameter settings. This would require detailed exploration of the hyperparameter space and will be subject to future research.

### Object classification properties

We showed that a binary SVM classifier optimized using higher-level representations performed better than a classifier trained on lower-level representations. This effect disappears when there is no regularization penalty (data not shown). Regularization of activity and synaptic strength forces the network to generate representations in which most neurons are inactive (or less active) and active neurons capture most of the information in the presented stimuli. This results in a representational code that allows better discrimination between object classes. Thus, regularization helps improve the accuracy of the classifiers based on representations in each area significantly above chance level. In combination with increasing feature complexity in the network, this leads to a modest but systematic increase in classification performance from lower to higher-levels in the network.

### Comparison with previous models

Most of the previously proposed predictive coding models utilized specific architectures targeting simulation of particular physiological phenomena (e.g. mismatch negativity (Wacongne et al., 2012)) or neuronal response properties (e.g. of V1 neurons (Rao and Ballard, 1999; Spratling, 2010)). (Rao and Ballard, 1999) proposed one of the first neural network models of predictive coding that was designed to study receptive field properties of V1 neurons such as Gabor filtering and end-stopping. With respect to their network, the specific advance of the current study is that it provides a methodology for building scalable, deep neural network models, e.g. to study neuronal properties of higher cortical areas. (Spratling, 2008) showed that predictive coding models can reproduce various effects associated with attention-like competition between spatial locations or stimulus features for processing. This study employed a network with two cortical regions, each having two to four neurons. A different study (Spratling, 2010) showed that predictive coding models can reproduce response properties of V1 neurons like orientation selectivity. These models consisted of a single cortical region corresponding to V1 and hence a top-down input was lacking. Both studies employed models with predefined synaptic strengths. In contrast, DHPC networks employ a Hebbian rule for adjusting synaptic strengths and estimating representations. They can be trained using images of essentially arbitrary dimensions. Further, DHPC networks not only showed basic properties like orientation selectivity at lower levels but simultaneously showed high stimulus selectivity and sparseness in higher areas, thus unifying these different phenomena in a single model.

(Spratling, 2012b) presented a predictive coding model in which synaptic strengths were adapted using rules that utilized locally available information. This study used models having one or two areas with specific, pre-set architectural parameters like receptive field size and size of image patches. Using predictive coding (Wacongne et al., 2012) showed that a network model trained to perform an oddball paradigm can reproduce different physiological properties associated with mismatch negativity. This study simulated a network architecture with two cortical columns, each of which had a pre-established selectivity for specific auditory tones. Unlike these studies (Spratling, 2012b; Wacongne et al., 2012), DHPC networks provide a mechanistic framework for developing predictive processing models with scalable architectural attributes corresponding to biological analogues like receptive field size and number of brain areas. In the current study, DHPC networks were scaled up to contain millions of synapses and thousands of neurons whereas most existing predictive coding models have simulated networks with up to hundreds of neurons and thousands of synapses. Furthermore, DHPC networks reproduce in the same architecture many attributes of neuronal responses without explicit a priori incorporation of these properties in the model. Probably, the approach closest to our work is by (Lotter et al., 2017) who employed networks consisting of stacked modules. This network was specifically designed to predict the next frame in videos and was trained using end-to-end error-backpropagation which is unlikely to be realized in the brain. However, an interesting aspect of this model is the use of recurrent representational units which allows the network to capture temporal dynamics of the input. This aspect will be an interesting direction of future research for the unsupervised Hebb-based models we proposed here.

### Anatomical substrate of predictive coding

An intriguing question related to predictive coding is its potential neuroanatomical substrate in the brain. Several studies have looked at possible biological realizations of predictive coding based on physiological and anatomical evidence (Bastos et al., 2012; Keller & Mrsic-Flogel, 2018; Pennartz, et al., 2019). DHPC networks are well compatible with insights from several experimental studies on predictive coding and error signalling (Leinweber et al., 2017; Schwiedrzik and Freiwald, 2017) and cortical connectivity (Douglas and Martin, 2004; Rockland and Pandya, 1979). However, some aspects of predictive coding that were highlighted by experimental studies have not yet been explicitly modeled by the current DHPC networks. A combination of experimental and modelling studies predicts that neurons coding inferential representations are present in superficial as well as deep layers of sensory cortical areas (Pennartz et al., 2019). Representation neurons in the deep layers are proposed to transmit top-down predictions to error neurons located in the superficial layers of the lower area they project to (Bastos et al., 2012; Pennartz et al., 2019). These error neurons also receive input from local representation neurons in superficial layers of the same area and transmit bottom-up errors to the granular layer of the higher area they project to. This anatomical configuration was not considered in the current DHPC networks because it requires explicitly modeling various cell types located in different neocortical layers and the interactions between them. This will be a direction of future research as it will help bridge the gap between theoretical models and biologically relevant aspects of cortical architectures implementing predictive coding.

## Acknowledgements

We would like to thank Walter Senn and Mihai Petrovici for helpful discussions and Sandra Diaz, Anna Lührs, Thomas Lippert for the use of supercomputers at the Jülich Supercomputing Centre, Forschungscentrum Jülich. Additionally, we are grateful to Surfsara for use of the Lisa cluster. This work was supported by the European Union’s Horizon 2020 Framework Programme for Research and Innovation under the Specific Grant Agreement No. 785907 (Human Brain Project SGA2 to C.M.A.P.).

---

**Figure S1.**
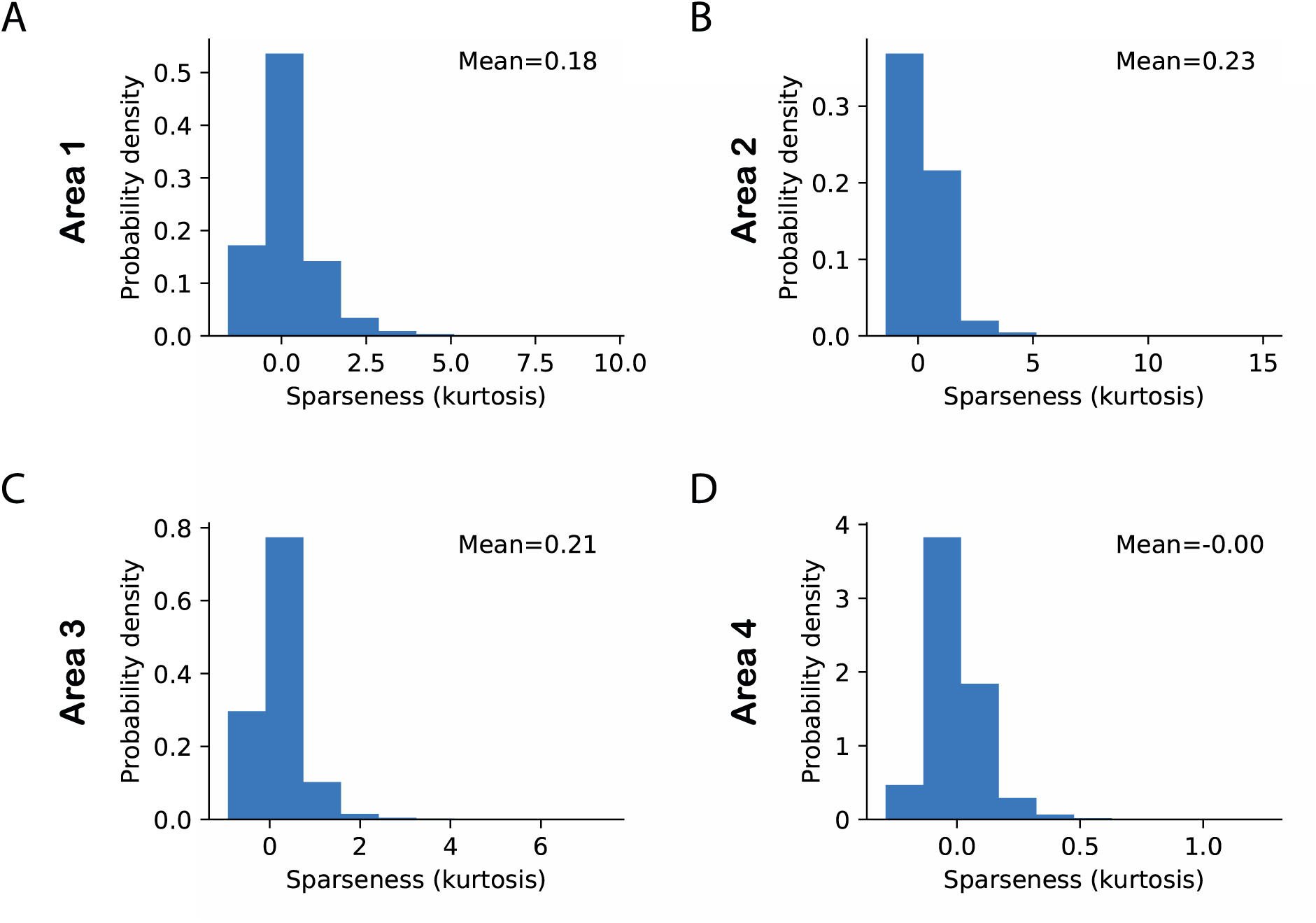
Sparseness in neuronal activity across ascending areas in a linear model without regularization of weights and activity. Sparseness was measured as the kurtosis across all neuronal responses in a given area and given a single stimulus. The mean value of sparseness (top right corner) was computed by averaging these estimates of kurtosis across all stimuli. (A-D) Distribution of sparseness in each area. We used models with a linear activation function as exemplars of models without regularization because ReLu enforces neural activity to be always positive, thereby requiring a strong regularization penalty. In the absence of regularization, the average sparseness in the model increased modestly from areas 1 and 2 and then decreased in areas 3 and 4. Despite its modest effect size, this pattern was observed across multiple models with a varying number of areas. This is attributed to the network property that all areas in the model (except the top area) infer causes that reconcile bottom-up and top-down information (Equation 4 and 6) whereas causes in the top area are only determined by bottom-up information. The lower constraint on the top area leads to lower sparseness in this area. This effect was not limited to the top area alone; it was generally applicable to areas in the hierarchy that were farther away from the sensory input layer.

**Figure S2.**
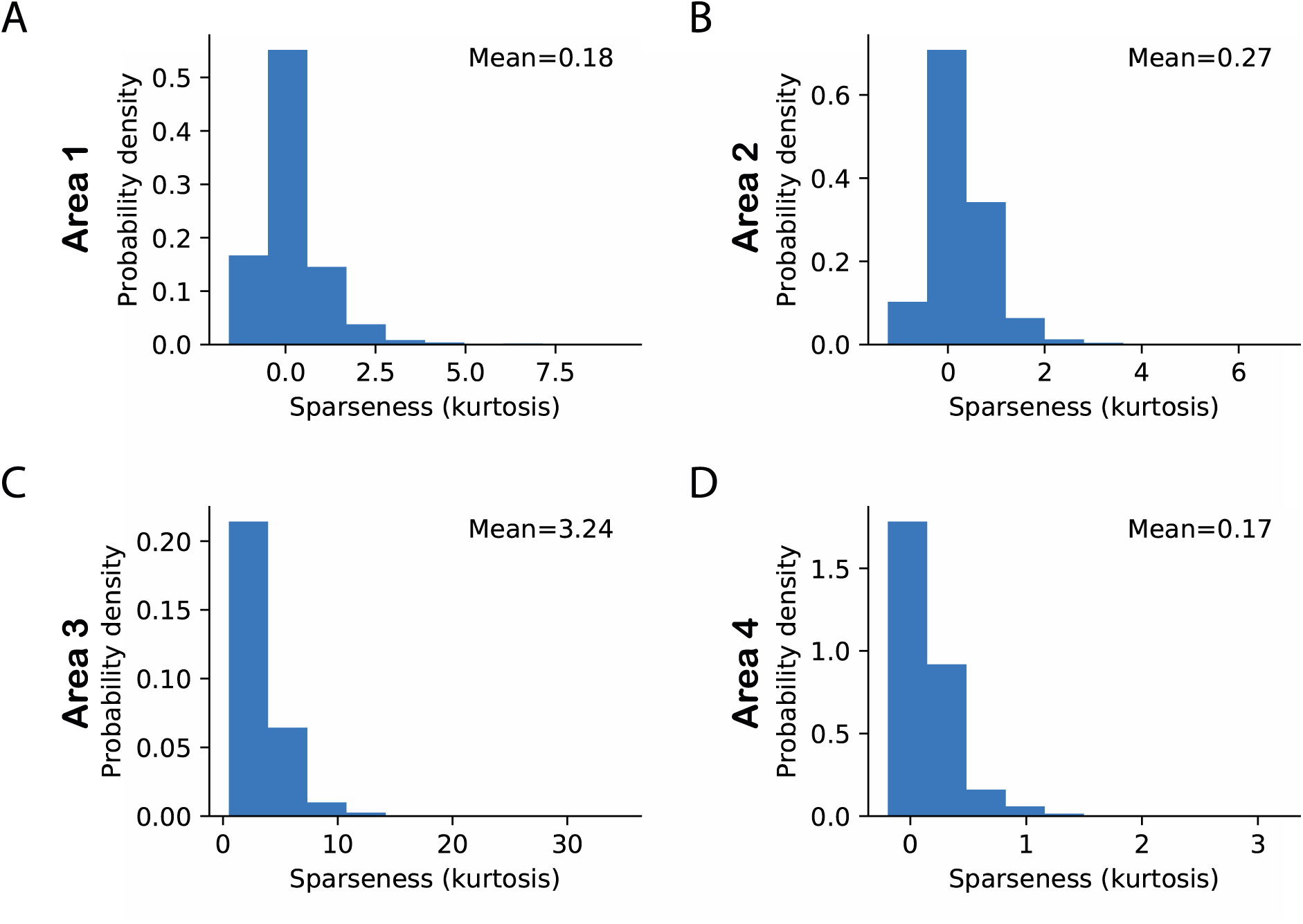
Sparseness in neuronal activity across ascending areas in a linear model with regularization only in the top area. Sparseness was quantified as in fig. S1. The mean sparseness (top right corner) was computed by averaging these estimates of kurtosis across all stimuli. (A-D) Distribution of sparseness in each area. Having regularization only in the top area presents an interesting case because this indirectly regularizes all other model areas. Regularization-induced sparseness in area 4 results in sparse top-down predictions propagating to area 3, which indirectly induces sparseness in area 3 representations. Compared to Figure S1, regularization results in an increase in sparseness in area 4 and indirectly leads to an increase in sparseness in areas lower than area 4. This effect is stronger in area 3 and becomes weaker as one moves away from the top area.

**Figure S3.**
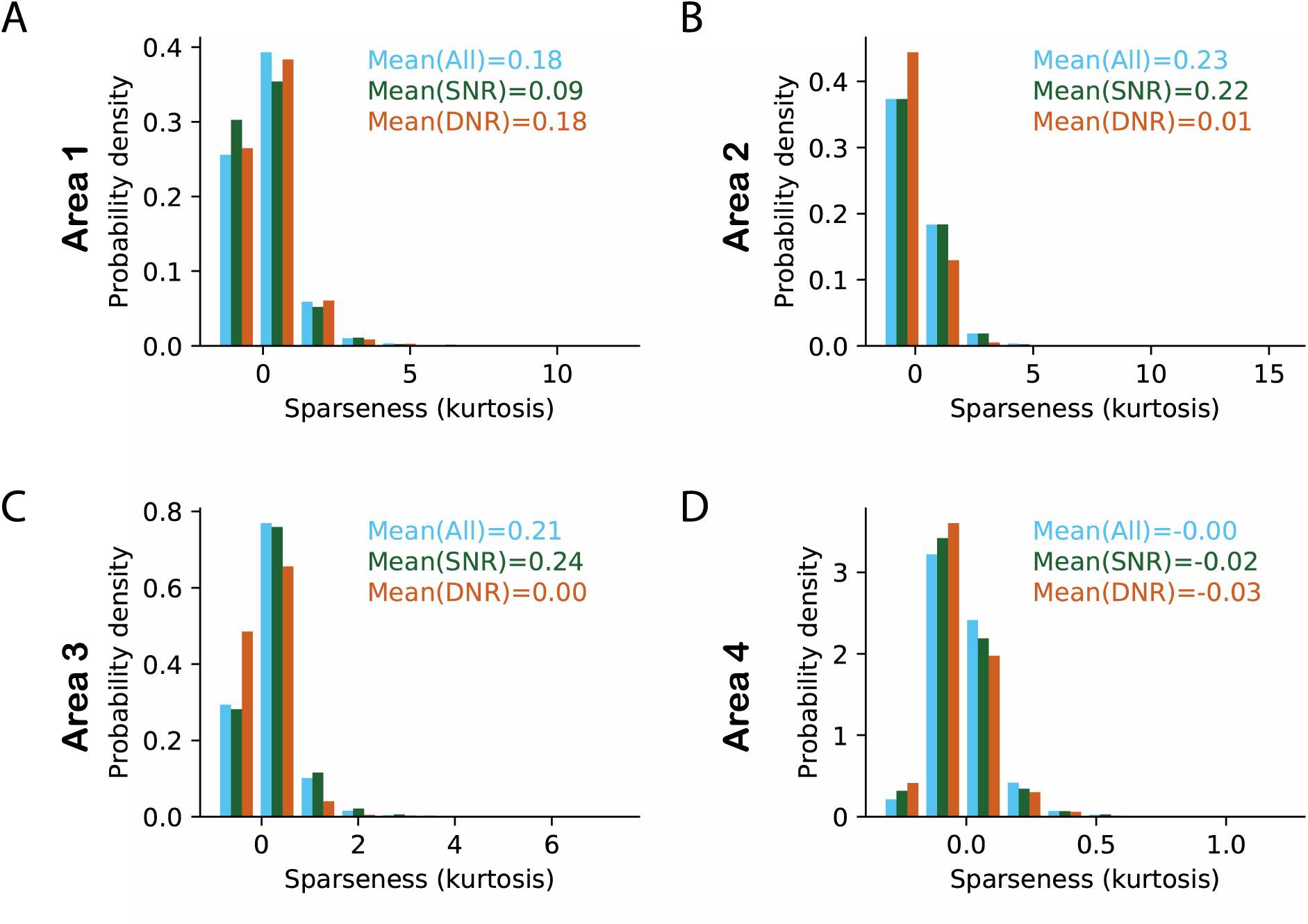
Effect of high selectivity and high dynamic response range neurons on sparseness in a linear model with no regularization. (A-D) Histogram of sparseness for three different populations of neurons. The distribution of sparseness for all neurons has been shown in blue. The population in which the top 10% of most selective neurons was removed (SNR) is shown in dark green and light brown color denotes the populations in which neurons with high dynamic response range were removed (DNR). Values in top right corner represent mean sparseness estimates for the different populations in corresponding colors. It can be observed that high-selectivity neurons contribute to sparseness in the lowest area (area 1) whereas in areas 2 and 3 the high dynamic range neurons contribute to sparseness. Despite modest effect sizes, this pattern was observed across multiple model variants. The effects are attributed to the network property that area 1 receives a bottom-up input based on a fixed visual image. Other areas in the network receive a bottom-up drive based on a constantly evolving set of latent representations. This leads to higher dynamic ranges in areas 2 to 3 and, as a result, sparseness is strongly determined by the dynamic response range in these areas.

**Figure S4.**
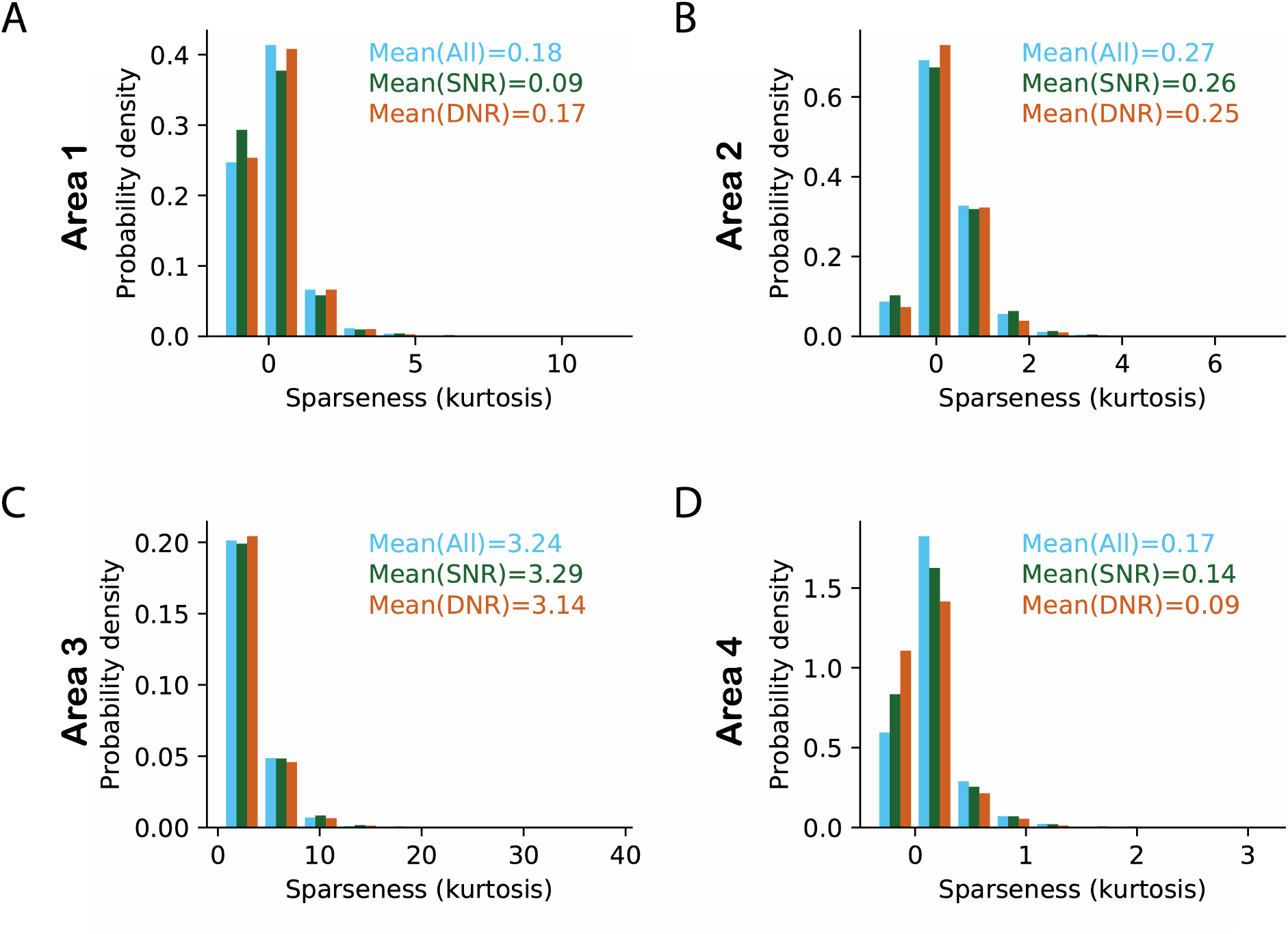
Effect of high selectivity and high dynamic response range neurons on sparseness in a linear model with regularization only in the top area. (A-D) Histograms of sparseness for three different populations of neurons. The distribution of sparseness for all neurons is shown in blue. For plotting conventions, see figure S3. As a result of adding regularization to the top area, the contribution of high dynamic range neurons to sparseness is weakened in areas 2 and 3 (cf. Figure S3). This effect likely arises because regularization, by definition, reduces neuronal activity; via a top-down spreading effect this leads to lower dynamic ranges in areas 2 and 3. In turn, this reduces the contribution of high dynamic range neurons to sparseness in these areas.

